## Supplementary Information for "Differences in structure and hibernation mechanism highlight diversification of the microsporidian ribosome"

**for**

**S1 Fig. Cryo-EM data collection and processing scheme.**

**S2 Fig. Global and local resolution estimation and visualization of the model-density fit.**

**S3 Fig. Conservation of Lso2 in eukaryotes and its ribosome interaction surfaces.**

**S4 Fig. Stepwise reduction of rRNA elements in microsporidia.**

**S1 Table. Cryo-EM data collection, refinement and model statistics.**

**S2 Table. Model composition & sequence information.**

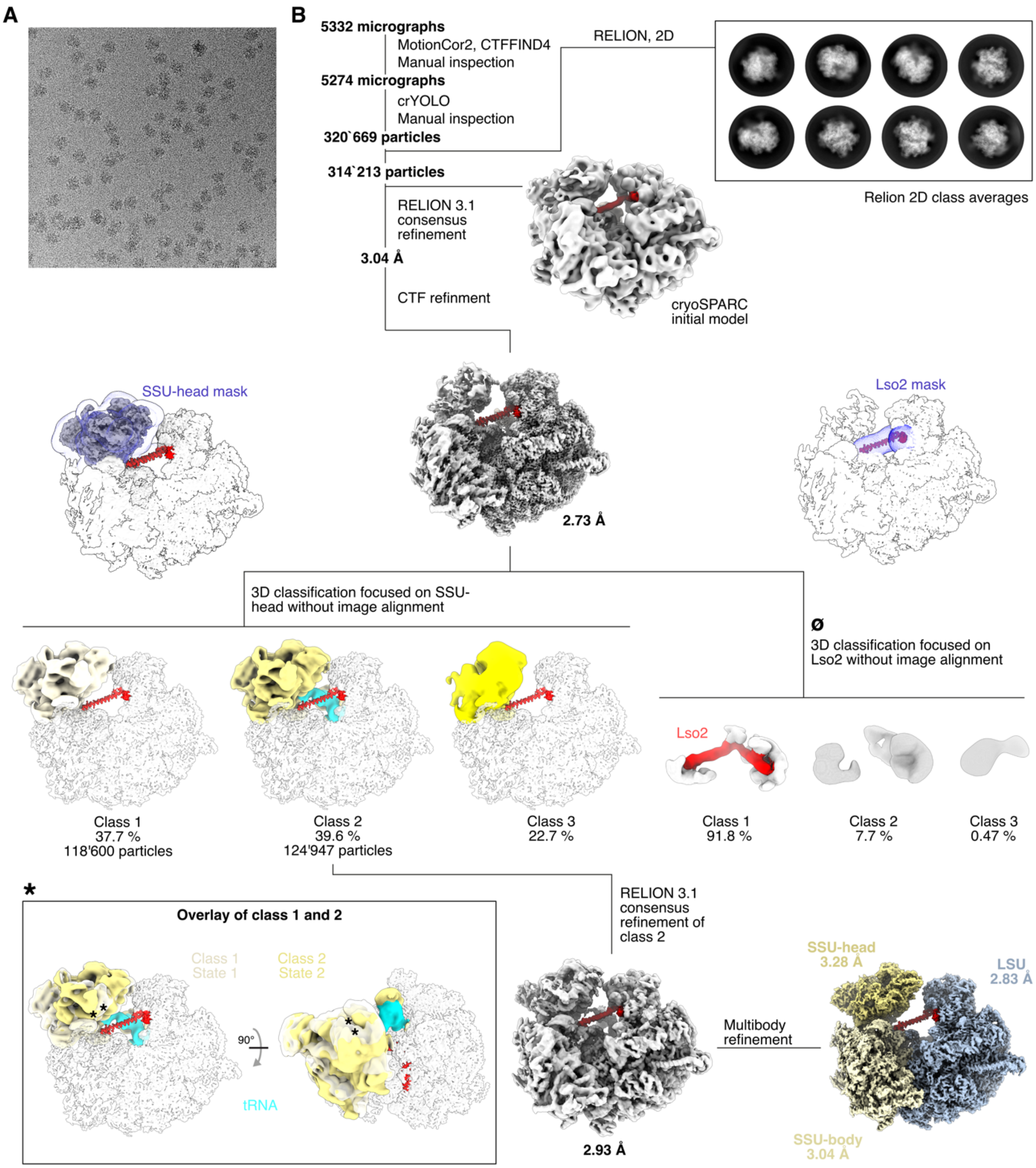


**S1 Fig. Cryo-EM data collection and processing scheme.** (**A**) Representative cryo-EM micrograph of the microsporidian ribosome. (**B**) The 5332 collected micrographs were manually inspected to remove those with drift, poor CTF fits, or low-quality ice, resulting in a total of 5274 micrographs. Particles were picked using crYOLO [1] and subjected to one round of 2D classification (representative 2D class averages shown) in RELION-3.1 [2] to remove picking contaminations. The initial model was generated using cryoSPARC [3]. A consensus refinement yielded a map at 3.0 Å resolution, which was improved further by per-particle CTF refinement to a resolution of 2.73 Å. To isolate the most populated conformation of the dynamic SSU-head region, a focused 3D classification was performed without image alignment. The resulting 3 classes of the SSU-head domain (different shades of yellow) are shown superimposed with the full consensus refined ribosome. Lso2 is highlighted in red. The class with the best resolved SSU-head, Class 2, contained additional density for an E-site tRNA (sky-blue), and was refined to an overall resolution of 2.93 Å. Multibody refinement yielded maps with resolutions of 3.28 Å for the SSU-head (EMD-11437-additional map 1), 3.04 Å for the SSU-body (EMD-11437-additional map 2), and 2.83 Å for the LSU (EMD-11437-additional map 3). These maps were combined using PHENIX combine-focused-maps (EMD-11437). (*****) The inset depicts a superposition of Class 1 and 2 to visualize the two conformational states of the SSU-head. (**ø**) To estimate the percentage of ribosomes bound to Lso2, a mask enclosing this region was used for a 3D classification without image alignment. Class 1 shows clear density for Lso2 suggesting that 91.8% of all ribosomes are bound by this factor.

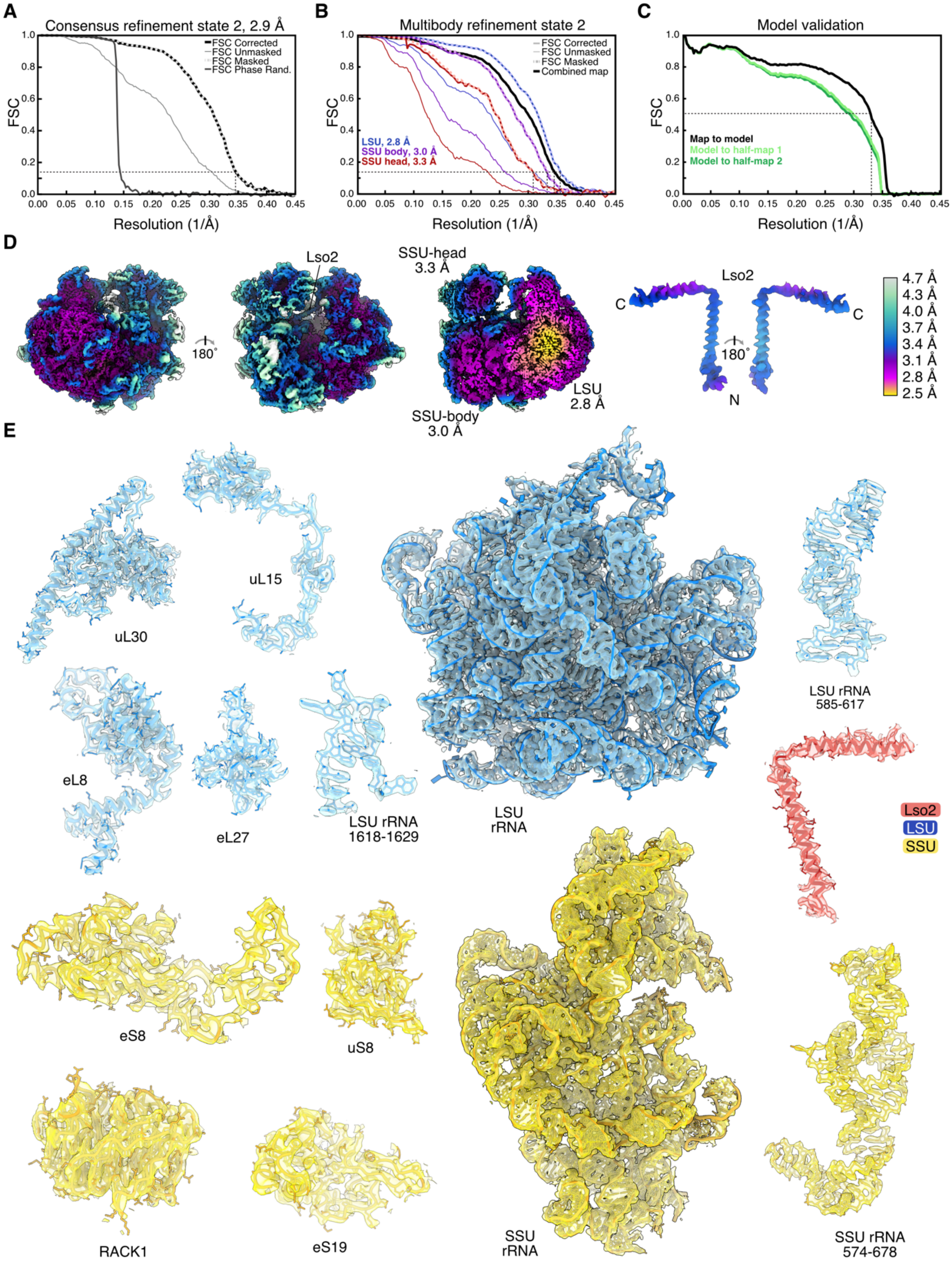


**S2 Fig. Global and local resolution estimation, model validation, and visualization of the model-density fit. (A-C)** Fourier Shell Correlation (FSC) curves of the consensus refined state 2 (A), the multibody refined maps and the combined final volume (B) and map-to model cross-validation (C). The thin-dashed line indicates an FSC value at 0.143 or 0.5. FSC. Curves were obtained from RELION-3.1 [2] (A,B) or EMAN2 [4] (C). (**D**) The final focused refined map is shown (left) next to a core-region cross-section (middle). Lso2 is also displayed in isolation (right). All maps are colored according to local resolution. Local resolution was estimated using RELION-3.1 and visualized in UCSF ChimeraX [5]. (**E**) Selected representative cryo-EM densities superimposed with the corresponding models, colored in blue (LSU), yellow (SSU) or red (Lso2).

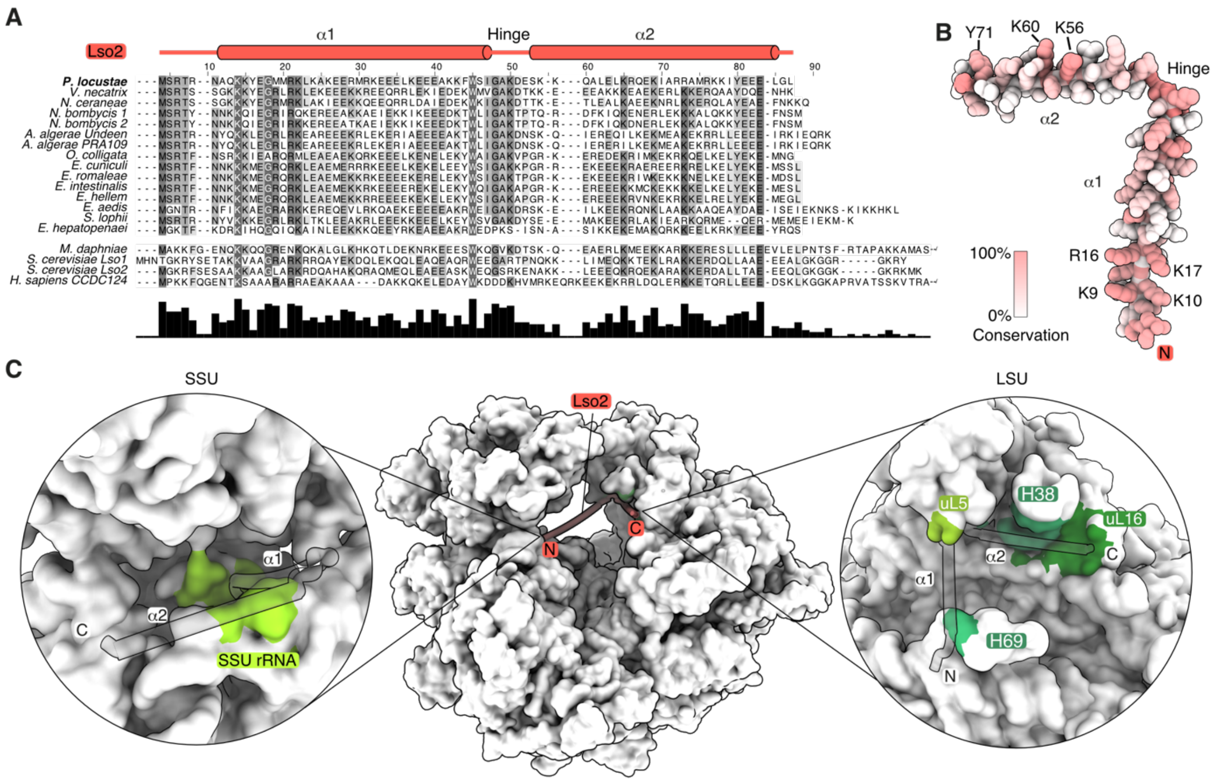


**S3 Fig. Conservation of Lso2 in eukaryotes and its ribosome interaction surfaces.** (**A**) A multiple sequence alignment of Lso2 from microsporidia and selected eukaryotes. The related *S. cerevisae* Lso1 is also included. The C-terminal ends of *M. daphnia* and *H. sapiens* have been truncated. The domain architecture of Lso2 is presented on the top. (**B**) Lso2 shown in isolation with side-chains as spheres, colored according to conservation from white (variable) to red (conserved). The conservation was calculated with HOMOLMAPPER [6], using the microsporidian sequences shown in (a). (**C**) Lso2-ribosome interaction interfaces (shade of green) were obtained using EBI PISA [7].

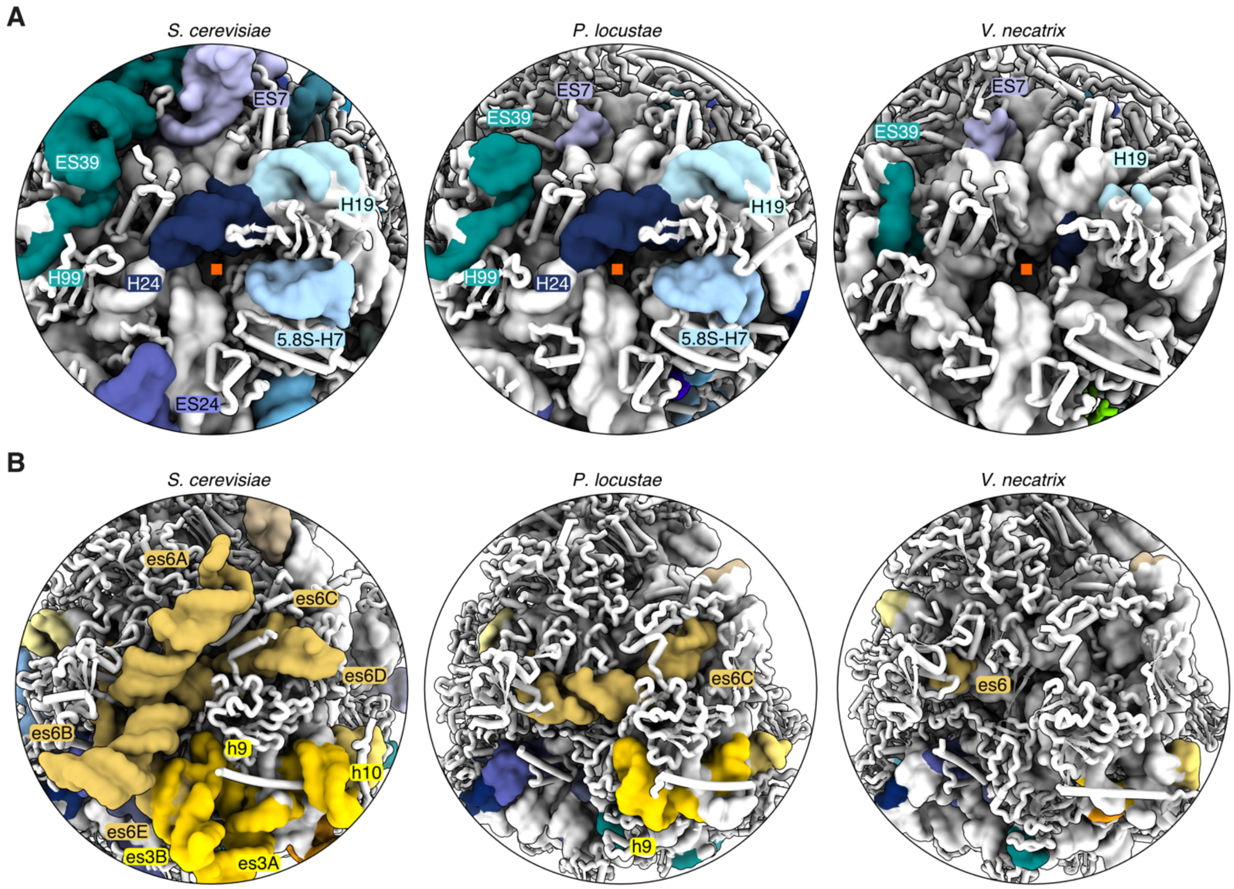


**S4 Fig. Stepwise reduction of rRNA elements in microsporidia.** (**A**) LSU region around the polypeptide exit tunnel, shown for *S. cerevisiae* (PDB 4V88, [8]), *P. locustae* (solved here), and *V. necatrix* (PDB 6RM3, [9]). Eukaryotic expansion segments and rRNA helices diminish from left to right. Peptide exit tunnels are denoted by a red square. (**B**) Reduction of the SSU expansion segments es6 and es3. *P. locustae* again represents an intermediate state in this reduction.

### Tables

**S1 Table**. **Cryo-EM data collection, refinement and model statistics.**

| | Combined focused maps | LSU | SSU-body | SSU-head |
|---|---|---|---|---|
| | EMD-11437<br><br>PDB-6ZU5 | EMD-11437-additional map 1 | EMD-11437-additional map 2 | EMD-11437-additional map 3 |
| **Data collection and processing** | | | | |
| Voltage (kV) | 300 | 300 | 300 | 300 |
| Pixel Size (Å) | 1.041 | 1.041 | 1.041 | 1.041 |
| Electron exposure (e-/Å2) | 28.6 | 28.6 | 28.6 | 28.6 |
| Defocus range (um) | 0.7-2um | 0.7-2um | 0.7-2um | 0.7-2um |
| Frames | 40 | 40 | 40 | 40 |
| Symmetry imposed | C1 | C1 | C1 | C1 |
| Initial particle images | 320,669 | 320,669 | 320,669 | 320,669 |
| Final particle images | 124,428 | 124,428 | 124,428 | 124,428 |
| Resolution (Å) | 2.9 | 2.83 | 3.04 | 3.28 |
| FSC threshold | 0.143 | 0.143 | 0.143 | 0.143 |
| Map sharpening B-Factor (Å2) | Combined map | -25.0655 | -21.2595 | -21.3032 |
| **Refinement** | | | | |
| Initial model used | 4V88 | | | |
| Model composition | | | | |
| Non hydrogen Atoms | 165,263 | | | |
| Protein residues | 10,308 | | | |
| RNA bases | 3,914 | | | |
| Ligands | 188 | | | |
| R.m.s deviations | | | | |
| Bond length (Å) | 0.010 | | | |
| Angles (°) | 0.897 | | | |
| Validation | | | | |
| MolProbity score | 1.47 | | | |
| Clashscore | 5.14 | | | |
| Poor rotamers (%) | 0.65 | | | |
| Good sugar puckers (%) | 99.11 | | | |
| Ramachandran | | | | |
| Favored (%) | 96.81 | | | |
| Allowed (%) | 3.11 | | | |
| Outliers (%) | 0.08 | | | |

**S2 Table. Model composition & sequence information.** Bolded and underlined sequences were modelled with side-chains while green regions were trimmed but still contain side-chain information. Sections indicated in yellow were modelled with poly-alanine stuctural elements, and the ubiquitin-moiety of eL40 is indicated in blue.

| Chain | Subunit | Sequence |
|---|---|---|
| S60 | 18S rRNA | C**ACCAGGUUGAUUCUGCCUGACGUGUGCGCUAGUCUUGGGGACUUAGCCAUGCAAGCCAGCGAAGC**GGC**GCGGAGCGGCGGACGGCUCAGUAGGACGGCGAUAAUCCACCCGCAGCAGAGACAAGGAUAACCGCGGCAAGCUGCGGCUAAGACAUGGACAGGCUGCGGCGCGGGGAGGCGCGGGCAGCGGACAGGGAGCGGCUGCGGCGGAGUUGCUGGCCCAUCAGCUGGUAGGUAGGGUAAGGGCCUACCUAGGCGACGACGGGUGACGGGGGGUGAGAGCCCGGAACCGGAGAGGGAGCCUGAGAAACAGCUCCCACGUCCAAGGACGGCAGCAGGCGCGGAACUUGCCCAAUCCCGGCGGGGGAGGCAGCCACAAGACGU**GGGGGCCCGGAA**CGCACGGCAAAGGAGUG**CGGGGCG**ACUGGAGGGCAAGUCUGGUGCCAGCAGCCGCG**GUA**AUUCCAGCUCCAGGAGCGCACAUGAGUGUUGCUGCAGUUAAAACGUCCGUAGUCGGCCCGGGCGCGCUGUGAGGAAACCGCGGGGCCCAAGGCGGCGGGCAGCACGGGAGCGGGGGCGACAGGAACGGACAACGGCCGCGGGAUCCGGGGGCGAGAGGUGAAAAUCGGGGACCCCGCGGGGACGGGCGGAAGCGAAGGCGGCGGCCGGGGACGUGUCCGUUGAUCAAGGACGAAGGCCGGAGGAUCAAAGAUGAUUAGAUACCGUCGUAGUUCCGGCAGUAAACGAUGCCGACGGGGCGAGGCCAGGGGGCUGUGUCCGGGAGAAAUUGGAGUGUUUGGGCUCUGGGGAUAGUACGGUCGCAAGACAGAAACUUAAAGAAAUUGACGGAAGGACACCACAAGGAGUGGAGUGUGCGGCUUAAUUUGACUCAACGCGGGGCAGCUUACCAGGGCCGGAUGC**GCGGGAGAUUGGCGGCGGAGCGCCCCCGAGAUUGC**GCAGGGAGUGGUGCAUGGCCGUUUGCAACACGUGGGGUGACCUGUCUGGUUGAUUCCGACAACGCGUGAGGCCUGGGGUGCAGCAGGGGCUGUGCGACGGGCGGCGGCAAGCUGCAGGAGGGCAGGCGAAAACAGGUCAGUGAUGCCCUCAGAUGCUCUGGGCUGCACGCGCACUACAAUGGCGCGGCGAGCGGGAGUGCCGGGAGGCGCGGGCGAGGCCGUGGCGCAGCAGGGAUGGGGGCCUGGAAGGGUGCCCUGAACGAGGAAUUCCUAGUAGCCGCGGGUCACCAAGCCGCGGCGAAUGAGUCCCUGUUCUUUGUACACACCGCCCGUCACUACCUAAGAUGGAUGUGCAGGCGAGGAGGCUGGGCAGGCACCCGAGCCUGUGCAACUAGAUAAGGUAUAAGUCGUAACAAGGCUGCAGUAGGUGAACCUGCGGCAGGAUCACUA** |
| L50 | 25S rRNA | **ACACACCCCGGUGGGGGAUCCCUCGGCCUGCGCGCCGGGCAAGGACGCGGACGCACGCGAUAGACGGCACGAUCCUCAGACACGACUGCCGGUCUCCGACAGCGGCGCAGCCGCAGACAACCCCCCGGACUUAAGCAUAUCACUAGGGGGCGGAGAAGAAACCAACAGGGAUUCCUGCAGUAGCGGCGAGCGAACAGGGACGAGCCCGCAUGGCAAUCGGCAUCGCCGAGUUGUGACAGCGCACCGCGAACGCCCCGGACAGGGCGGCCACAGAGGGCGACAGCCCCGUAGCAGCGCGCAGCGGAGCGAGUAGCGCUGCUUGGUCAUGCAGCGCGAAGCGGUGGUGGCGCCAUCGAAGGCUAAAUACGCCGCAGGACCGAUAGCGCACAAGUACCGCGAGGGGACGGCGACGAGCAGCCCGCAGGGGCGGCGAAAGCGUGAAACCACCGGGGCGCCCACUUGUGGGCCCCGUCUUGAAACACGGACCAAGGAGUGCAUGUGCGCAGCGAGUCCGCUCCGCGGCGCAGCGAAGGCCAUCGAGCUGCGCACAUGCGACCCGAUAGGCAGUGAACUACGCCUGGGCAGGGCGAAGCCCGCGGAAACGCAGGUGGAGGCCCCGAGCCGUUCUGACGUGCAAUUCGAUGGCGCGACCUGGGCGUAGCGGCGAAAGACCAAUCGAACUGCCUGGUAGCUGGUUCCCUCCGAAAUGUCCCGCAGGACAGCGGGCGCCCCGCAGGUCUGCCGCGUAGAGCAAUGGCGCGGCGUCCGG**CAG**CGCCGGCGCACCCCCAAACUGCGAAGCGGCAGGGCGCGCGCAGCAGCGUGCGCGCGCACAACUGCGGGCGCCUAGUGGGCCGCCGCUGGUAAGCAGCGCCGGCAAUGAGGACACAACCUCGUGCGCGGGCAAGGGACCCCAGCUGCGCACACAGACGAAGGGCGCGGGCGC**GUCGCGACAGCAGGGCGGUGGCCAUAGAGGUCGGCACCCGCUAAGAACCGUGUUGCAACGUACCUGCCGAACAC**GCCCGCCCCGAAAAUGGACGGUGCUCAGCGCAGCCCCGACCCCGCGCACGCACAGCGUGGUAGGAGGGCGCGCCGGCGCCGCAGAAGCGCAUGCGUGCGCAUGCGUGGAGGCACCCGCGGCGCAGAUCUUGGUGGCAGUAGCACACUCGGGCGCGAGCCCCGAGGGCCGGGAGACGGGUUCUUCCGCCAGGCCGCUCCGCGGAAGGUGAGCCGGGUCCUAAGGACGCGCUGGCCCGCAACCGACAGGCAAGCGGGCACACAUUCCCGCGCCGUGUGCCAUGCGGCAACGCACCGUGCGCGGCCGGGCGCAGGGCUGGCGCCGGGGGCCCUCCUCCCCCG**CAAA**GCGGCCCGCCUGCGGACUCUUGCAGCACGAGGCAGCCCGCGCCGC**GUGGCGGGGCCGUCGCCGCG**CGCCAGGACUCGCCCCCCGUGAAGCCCCGCGCACGCACACACACGCCCGUACCAAUCCGCACCAGGGCUCCAGGGCGCGCACCCCACGGCCAGGGCCCACGCAGGUUUGGGAAUUCGGCAAGCUGGAUCCGCAACCUCGGGACAAGGAUUGGCUCCGGGCGCCGGAGCUGUCGCUUCCAAGGGGAAUCCGACUGUUUAGUAAAAACAUAGCCUUGCGCCGCACGCAAGGUGAAUUCUGCCCAGUGCCCGG**GAC**GUCACGCCGGCGCGACCCGCGCACGCACGGGUCAACGGCGGGAGUAACUAUGACUCUCUUAAGGUAGCCAAACGCCUCGUCAUCUAAUUAGUGACGCGCAUGAAUGGAGCAACGAGAUUCCCACUGUCCCUACCUGCUCCCCAGCGAACCCACUGCCAAGGGAACGGGCUUGGCGCAGUCAGCGGGGAAAGAAGACCCUGUUGAGCUUGACUCUAGUGUGGGGCCGCGGC**GCGCCGCGCCGGCGUAGGCAGGUGGGAGGUGCGCCGUGAGUGAAAGACCACUGCGCGCG**CGCGCCCGCUUCGCGCAGCAACGCCCCCAGAUGGGGAGUUUGGCUGGGGCGGCACGUCUGCUAGACCCCAACGCAGACGUCCUACGGUGGGCUCAGCGCGGACAGAACCCGCGCGUCGAGCACAAGGGCAAACGCCCGCCUCACGGCGCCCCCCGGGUGCCGGCGGGAAACCGGGGCCUAGCGAUCCCUCGCGCAUGCACGCCGCGUCGCGGGGGUGGCUGAAAAGUUACCACAGGGAUAACUGGCUUGUGGCGGCCAAGCGUCCGCAGCGACGCCGCUUUUUGAUUCUUCGAUGUCGGCUCUUCCUAGCAUGGCGUGGCAGCGCGCGCCAAGUGUUGGAUUGUUCACCCACUGACAGGGAACGUGAGCUGGGUUUAGACCGUCGUGAGACAGGUUAGUUUUACCCUACUGAGC**GC**GGACACACCGGGCAGCGCGGGCUAGUACGAGAGGAACGCCCGUGCGGGGCCGCUGGUCCGCGCCUGUCCGACAGGGCAGGUGCGCCGCUACGCCCCGUGCGUGUACGGCUGGAC**GC**CUCUAAGCCGGAGCCGCCCCCCCGUGUGUCUAAAC**CCCUG**GUUUCCGCCCCCCGCGACCACGACGCGGCCGGGGGCUGGUGCUGUGCGCGUGCGAGCUCUGCGAGCCGCUGAGGCUUCCAGACCCCUGCGGGGUGUU** |
| L70 | 5S rRNA | **AGCUGCGGUCAUAUCCACAGUAGAGCACCAGAACCCGUCAGAUCUCUGAAGUUAAGCCUGUGAGAGCCUGUGGAGUACUUGGGUGGGCGACCACCUGGGAAAUGCAGGUACUGUAGCUU** |
| LNN | Lso2 | M**SRTRNAQKKYEGMMRKLKAKEERMRKEEELKEEEAKKFWSIGAKDESKKQALELKRQEKIARRAMRKKIYEEELG**L |
| SA0 | uS2 | MQKEIEIPEDFVKAIILSNAHLGSTTATTNFKQYIYGRRKNDLIHVIDINEMWKKLIIAARLFVSIEDPSSIIVASNKKFGKRAVLRFCSETGATPLIGRFIPGGFTNQVINDVKESRLVVVSDTYADKQTVMEASYVNTPCIAFCNSDNSLSRVDVVIPMNNRSPLSIGCGFYLFAKVIKYIKGEESFDVDVRSEIELYFYRDPVELDMIIQEQKEKNETSFQVKDAAPEVPAEAEQWADA |
| SB0 | eS1 | MAKKDQPVTKKSLKKGLKKKATD**SFAKKEWFSLRIPSGFVTAEPGKTLCTKQNTKANLEKYLLGRNFEVNQGDLNPGNEESAVRKFRFIVNNVNGRIAESEFNGMELTTEKKRGIVRKWHTLIEAFRDVCTKDGYHLRVFVTALTKHR**PGSVS**RTCYAKTSDVKKVRRLMFEVIENEVSGSELAKVVKKLSVEKLGKEIEKKASVILPLQNCYTAKVKVVKRPR**PEVIRN |
| SC0 | uS5 | MESSVAGRNTGPLPRKKEEAVRPRVPRQQDK**EWVPTTDLGILVKMGKINMEDIFQYSLKIKESGIVDHLLKGQLKEEVIKIKSVQKQTRAGQRTRMKVVVVVGNGAGYVGIGTKTSKEAATAIKGAIERAKCAIRPVRLGFWGNNYGKEHTVPCKATGKSGSVIMRVIPAPKGSGIRAGSVPKKIFQLAGVKDVFTLSKGQTSTTENFAKATVSALDKSSCFFVPEFWREKEPELNPLSRYSRLIKEYREKI**ARK |
| SD0 | uS3 | M**VSTKQVLSKFIEKGIMKAELNEFFSKEFEDEVYSGMEMKPHEIPVQITVRVNKTHEVIGERKSRLNRIKHVLMERFPLLRTGVEINVEMVKNKGLCPKTQAEYIKAKLIENVPFRRAVNSVMRSIKESGAQGCVIIVSGKLKGQRARSVKYADGLLIHTGSSKEDYVKEALSVVHLKQGVIGIKVRIMLPYDPENVNGPSKVISDKIKIFEPK**ELY |
| SE0 | eS4 | M**TRGPRRHLKRLNAPSSWQLDKLGGKYAPKPSSGPHRLSECIPLCILLTRKLEYATTTKELKYILRKKMIKVNGVVRTDKDFPVGFMDVISIDTTNENMRLLYSNNRAFMLHNISSEEATYRLCRVSKKKMGIGGIPYIYTQDGSTFRYCDPKINVNDVVKVNIKEKKVVGFLPFKSGMKVFITKGKNLGCIGVISGIEKHMGSHDIAYVVDAKGRSFSTRTSNAFIIGDSESYWISLPEGEGIKISELEKSNIKYGE**IASVPVAEE |
| SF0 | uS7 | PMSIACDSGE**LLLFDKYSRDEVVVNDRSLKNYIFLRSLSSYPHTASRKGSRQFGKGEIPIEERFVCGLMRRGRNSGKKRLAMAALRNAFIIINTLTGKNPLQVLVDALVNCGPREDCARIGKGGAMKRTSVDVSPLRRLNVAIFLVCKGIRQACMKSTKTIAEAIADELIAASRNSPNSFGVKKKDEIERMAKS**NR |
| SG0 | eS6 | **MKLNVAYPAKGTQKMIPIDFQTSMRLYGLRLGAQVEGELIGEDFRGYVFKITGGDDVSGLPMKDGVFTDKRVRLLLKKGNKGYRCREKGVRKRKSVRGCIISGEIAVVSMIILKEGEKP**IEGLTD**RVVECSHLPKRASKLRKLFNIPKDANIVEFIKEQIRNNSDSEKKIRYPRIKVTRLITPMRIERKRKRLEEKEARKIKSENERKEYMKKYYPEKI**ESL |
| SH0 | eS7 | MSEQTEHEKKVAQEIKKLYSQDMSGEFGDKLSDIRIQTIDREQGKKILLIRVPEEVLEVVHIGYSKLSGKLKALYPDFHTLVIRNTPIVNVSSIKGNVYRI**DSKNTKKSVHENWIRDLCYPALVEMRKTDVFNGCERIESALVSSNSNYSDDDFAAMECAFKTLTGRTIHYGNIFY** |
| SI0 | eS8 | M**GLSRSNRHKRTKTGAKKPIFCKKRKFHMARQPSNTKIGEDKLQELRCRGGVIKKRALRLSSGNFTCRTHGFSYKTKIMQVMYHPSSNELMRTNTLTKSAVIKLDSKQFKEKFDSILSTDSELKNTDPLFFSNLDAGKLYGVIVSRPGQCGRADGYILQGDELNFYINKFK**KKAR |
| SJ0 | uS4 | M**ARPKYSKRSRVPRKPYEKDRLIDEMKLLGEFGLKNKRELWTQEKICDDIKKRARDLLISTDESEIIINGRALLNKLIKLGIFSDVDFKDMDDIKRNLEKVLDLTASHFLRRRLQHRVFEACLAKSVHHARNLINHRHITVRGCVIDKPGYMVFAENEGHIEINK**FSSVGGAKPGRSKKKHAVAQGE |
| SK0 | eS10 | M**LIPTKNILAVKRHMLEKRCIVVEDTPKGNHPVLNIPNLQVLNIMKSLVSKRYVKKTFCWRHGYYFLE**DTAVDVLKDQLCMSDDEAAGLYAPSAVVSNIKAQ |
| SL0 | uS17 | MGDILDKA**KTYPKQKKPHL**VSRREGKDI**APERYYKNIGMGYNVPETAINGTYIDKKCPFTGNVTIRGRIFKGEVFKMKQEKTIVVVKRYFHYNKKYKRYERRNTKFNVHLSPCFFGLVRVGDTVTCGETRALSKTKHFAVIDYKQKDS**EDGQYAQFTGFS |
| SN0 | uS15 | M**ARMHSKGRGVSESTVPYNRVRPHWVTQSKDEVVGTILSLSRKGLKPSEIGKIMRDEYAIGHVRNVTGMNILQILRAHGLAPAIPEDLCALLDKRAGILSHLNTFRNDKSARYRLNLVESRMYRLARYYKRTMVLPATWKPHA**ASK |
| SO0 | uS11 | MTEDIS**TKDLVFGVAHILATSNDTFVHITDLTGSETVCKVSGGMKVKAHRDESSPYAAMLAAQDAAQKAIERGITAVHVKLRGRGGVKSAIPGPGAQAALRALIRNGLKLGRVEDATPTPTDRTRRKGGRRGRRL** |
| SP0 | uS19 | MEGDVEKERRQFK**KYTYRGVPLSELVKMPLLKFSELLTSSKRRRLRRGISERELDLLKKCALSKKNAEGTQEKPDMVKTQARTMVIVPQMVGCVIGVHNGRVYVPVEIKPEMIGYYLKDFSDTKT**CSTHGKPGVGATSGSKFVPLK |
| SW0 | uS8 | **MNRLAATCKAINNANRSGKRQVLIRFVTKTTRKFLEKMQVCGYISTLTYIDDHRQGKVIIGLNGRLNKCGAICPNYCLTLKDVEKFKDKILPARQFGHLILNTSKGMIDHNECVTMKSGGEVLGFFY** |
| SQ0 | uS9 | AIPM**AKKVIAIGQKKTAKARSECMEGDVFSVTVNNVPLRLLPDDMMASKLRDIITRVGLKYFENLVFSITVRGGGPVSKIYAVRQAFAKSLVAYYGTYMDEWTRQEIRSNLLSFDKFTLVADPRRAEPKKY**GGRGARARRQKSYR |
| SR0 | eS17 | MG**RVRNSNVKRAAKNIVEKYFPKLDCDFHHNKRVLESVTEIQTKRFKNQIAGYVTHLYTRVEKGKVKKIFI**KKHEEERERRENMIPSSSIMDTEKIEVDDITMQMIKDYCYRGNYVVSVSEEQ |
| SS0 | uS13 | MEGRKS**LFETFTSEPAQIQHIIRMLNTNIDGSRRVVHALTQIKGIGKRMSDAIVKRTGVDASKRMGALSASELELLQDGIQNPAKYGIPRWMFNHQFDAVDGTDSHLIGNQIDANLRFYIEKGKKMKHIRVCRLARGLKVRGQRTKSNG**RGGKTIGVSRKK |
| ST0 | eS19 | **MKELFMVRADALNEALAGYLKSKNIVKLPENFDIIKTSHGKQRAPSDKDWYYKRAASIFRKLLIAMHEEKRIGVKRLSRKYGCAKNRGSRPSKHVDGSRGHIRKIMQDLERAGLIVKDGNYRKVSDMGIIETGQLISQI**KE |
| SU0 | uS10 | MELEIDEKKY**SVDINLEGTSVEEIEKVCRDFYAYAKTLVPDMGGVKRLQTKKGKITTRRSPCGEGTNTWSRYRISVHRRSFSAHLYPSVLEKFATFLKGSNVEISLVIK**SN |
| SV0 | eS21 | MS**FEQRICTFTGRPIPHTDNASVQLTFV**ELDSDGRATDKV**LIYNVCGKIRKDGTIDEILTNTLAQ**N |

| | | |
|---|---|---|
| SX0 | uS12 | MKGLYCAKKLKRNRQAKRRNDPTYRKRLLGTKYKQDIIGVAPQAKGIVLEKIEVEAKQPNSAIRKAVRVQLIKNGKKVSAFVPYDGAINDIDMNDIVTLEGFGKKGRSKGDIPGIRYKVCKVQNVSLLALFRGKKDKPSR* |
| SY0 | eS24 | MSLETQICMMRENRPLQRREVVLSISHPKSRTPSKEDICSQISSLFKVRKELIIVSGCSTRFGTHQTKCKVRIYESPEMLREIERDFVVKRKTGEEVRKKKARRVRKKERKEKAKIFGTLRRHLKKAEKRAQK |
| SZ0 | eS25 | MVKKVLESKEKKAAKIASTSNKEKKKWTQGKTREAVRRSVTVEADVFSKIERDVAKASLVTAPSVAEKFNLNVGVAQKILEHLCAGGVLCCLSRNSRLRLYSRAQKVGARPADTTLPAAEAPAQTE |
| SAA | eS26 | MPVKRKNHGRAKKNRGHVKLIRCDNCASAVPKDKAIKRFQIKSLIEAAAHDDVRTATIYEEYEVPKFFHKNQYCVSCAVHLKAVRCRSAEGRKDRSNPHARVKQ |
| SBB | eS27 | MAIKDLAFPTAEELRTTCKRKRLIPQNNSYFLYLRCKSCDIIILAYSHSQTRRTCPGCNMVMLMPKGGKARIEGDVKIKKKIKRLVE |
| SCC | eS28 | MTDQVFLGEVMQLYNKTGPGGSITLCQLKLEDTGRMLHRAVIGPIKIGDVVTLLDCEREHRRGRF |
| SDD | uS14 | MEENIKIKPGSYAGRVDTRKFGRGSRSCRACFTHRGIIRQYNLYLCRRCFREYALEIGFKKVD |
| SEE | eS30 | LESKGSIARAGKVRNQTPKVEKQEKPKALTGRARKRALYERRLEANFFETRKMKMNPNFS |
| SGG | RACK1 | MTTTARLNEVATFSGHKDAVMALGTCETPDAKLLFSASRDRRVLGWSLGEGAMFGRIVKEFTKHGHSVNDVAVARSGSFVVSAGCDGLGRIIDVQSGERTLLRGHESDLTCAAINCQENKIVTGSVDKTLRLWNMCGELQHTFDAAIECAHEDWVMCAEFRPLDENEVVSGSVDGTVKVWDIDARVVKSTFFDGCLIQSCAEGSEYRTRPVADGSFAVRALALSADGSCCAYGGSNCKTYILNLAESEAIAAFETDTPVSALAFGLTDVILACGTQDKIYIWDVVSNCLLAVADLSAHGKRVRCRSLVWTTSNLIAGLGNGKILVFEFVR |
| LA0 | uL2 | MSKVIRRLRLLKPHHRPKAMWKVGETRIPVVSETMHGVVSEIVHERGKVAPLAKIRVDTGKCVRRELLVAVEGNYVGQKVEIGDSVPVAVGNALKLKNIPEGTGVCSVERRPYDGGKMAKSSGAYVTVVGHNRDTNITTVRLPSGEKRSVSSECRAVVGVIAGGGVNEKPLLKASRAHYRAKARGLYWPTVRGVAMNPVDHPHGGGNKQHIGHPSTISKHAPPGQKVGLVAARRTGLRRGSKKVLNK |
| LB0 | uL3 | MSCRKFEAPRHGSLAYMPRRRARSVKQSIRAFEKDNPEDPIHLTAFYVYKAGMTHVVRNKAMNDKKGTIKEVTESVTILEAPPMVVFGIVGYVNTPQGLKINKTLLSSHINESVLRRFYRKFYLSKKRMFSSARRKAGQKELDADILVLKDSDVIRVLAHTQVEKIKSIRTKKAHISEIQVNGGTVNDKVEWAVSMLEREVKISDVFSTNEFVDTIGVTKGKGFQGVTKRFGTRILPRKTNKGRRKVACIGAWHPANVLRTVPRAGQLGFHRRTELNKLIYLIGNGKEEIKTDFDPTLKSINPMGGFPHYGLVNNDFLMVKGGITGPVKRVLAIRKNLIGKKNNENIQIKFIDTSSKIGSGRFQTSEEKRAFFGITKKDVSEEIK |
| LC0 | uL4 | MRDTVNCYGIDGETVEKQLEMPDVLRVPIRKDLVEDAFRCVRMDNRQPYAVSPNAGMQHSAHSWGTGRAMARVPRVSGSGTTRSGQGAFANFCRKGRLAHPTKVIRRWQRKFNLNAKRHAEAMALAATAIPPLVESRGHRIAGVKMIPLVVSNSIKEIKSTKEAFEMLKRFGLAEELARVKESKCIRAGKGKMRNRRYVMKKGLLIIYDNQSDIQKAFRNIAGVDLACVDSLSLLDLCPGSHLGRLVMWTLGAFEKLNEIYGQYGKEAPLTSGYFLPTNVVSKDDVESLFFSDEIQAFLDVPNLIKYEKTSRKPETIESLNPYLNLMESN |
| LD0 | uL18 | MTDNKIKKSYFSRFQTKLRRRREGKTDYKHRYNLIRQDVNKHGLMKIRLVVRITNSRIICEILRAHVDGDRSIAYADSTELKRYGITFGLKNYTAAYATGLLVACRYNNKIAGEGPRPECYLDIGLRRSTRGARVFGAMKGALDGGLVMPHSLKRVPGYVSEEEFDSEVFRNKLFGKILAGYMKEMMENYPEKYKKTFQEYIKKGINPDDLENIYENAFKKIREDPSRVSKTHGDYSIFKEFKRVRLSKEERAARSRAKLLDIEVK |
| LE0 | eL6 | MIDTSKLSMLSRRGIKVIPEEAGLYPSDDLPAYIEKLEKRMQKKPRVRRTDLVKGCIVVVLEGDFTARRVVFLKQVENNKALCCGPAPINNVPFFVIDERYLLRTSTVLDLKEDVNIDISTVFESKRGVVADRMDIDANSDQKRIENAIVDAVSSIRFMKRYLATPFKMPKFKSVSSLKF |
| LF0 | uL30 | MDQEVPYSYRRKMEYEERMEVLSRKMNEEYQARREENARYALQRTKELIAEYKEHMKKEEDARREAEARGAFYVPKEPEFYVVVRIRGIHKVPPRERKILELIRLKKPNHAVFVRNNAPMRAMLHKVRAYIAFGFADIHLLRTLVYKRGMAKVSKRPLANNSSVFKIGQDMRLNLTNEAIEDHFGGRIRCVEELIYQIYMGTDLFKKANNFLWPFTLCSPRKGFGGRKAKDITQGGSTGCHYDKIGNLIYRMIE |
| LG0 | eL8 | MASQAKREMIERRRNPRDLDQKHQREERKALIAERVHLYRNALRIPPAVYQFSTFLNQQDMDQVLSLFRNYIPETRAEKKKRLMSENPRAGRKPILIKFGLKHVTDLIERKEARIVIIASDVDPIEVVVFLPTLCRKMGIPYAIVNGKKELGTLVHLKSTSCICLCDVAPKDSVEFKNILAKINAIFMDNYEITMKKWGGGALLRDKPQESSAPQA |
| LH0 | uL6 | MKRLLCEEKVEIPEGCSVEILERVMTVRGKRATAVRDLSHFVLTMDVHEGHVRLRLWNGTNRERSKLITCASVIRNCIVGCMSGYEYTLKVVYKHFPMSVAIEDDGKTVVVKNFLGQKHARRYKMRGDSIARLGTEKDTFVVEGSSLEDVSQSAGTIQENCQVKKIDSRTFLDGIYFLSRNVVGAK |
| LI0 | uL16 | MGRRPARCYRYCKNKPYPKSRFCRGVPDPKLAIYDLGRKRARVTEFPLSVHLVSNEREQLSAEALEAARIAANRYMIKHAGKDNFHMRIRVHPLHVIRINKMLSCAGADRLQTGMRGSFGKPYGRVARVDFGQEILSIRTKDAFKAVACEALRRAKAKFPGHQQIKVSCAFGFTGISREEFERLRKEGKLISQGSHVTIIKEKGSIVGYKEKLSKAIQ |
| LJ0 | uL5 | MTEQLENPMRQIRIEKLCINCCVGESGEKLNRALKVLEQISGQKPCTGKARLTIRGFGIRRNEKISAYVTVRGEKAREILNNALKVKEYEIKKSSFSNTGGFGFGIDEHIDLGIKYDPSIGIYGMDFIVVLSRPGLRVSKRRIKKSRVGNKQRISKEEAMQWFVKNYDGVLLNK |
| LL0 | eL13 | MKHNNTLPNNHFKKTAIRFKTWFNQPAKKEKRIQLRKEKAEKMYPMPVEKLRPIVRCSSIRYNIKQRLGRGFTPEECRAAGLDYNYARTIGIAVDMRRKNMNKETFDQNVERIKVYTSRLTFYKDRKEAKASGAAQHIGKIMPPKKTPPVVQTIKVEEIAQKFPTK |
| LO0 | uL13 | MINKEIIVDGTGHIAGKLGTYIAKKLLEGYTITVLCAESIVLTGPIHRTKLRYKDYLNKRCLVNPLRGPFHYKEPSKLFMRLVKRMVPYKKKRGAAALQRLQVFEGIPEKFENTERSICPRALLEYCANPIRKSATYGKLLSEFGWKHLNITEEMKKKVLQREEKAKKEKDARMEEIQRIRESSSFNKEVEEIMSRIE |
| LM0 | eL14 | MAKNFVQVGRLATPRFLAQRNILYVITDIQDDSMVIVQDAQGTRKLVSVKSLHLMDDVVEIHRDMSVEEVSKKIPQQKEEEVSENDFERFKRELRTEIEEEVLRSRGF |
| LN0 | eL15 | MSASEYLREIRKKKQSDLARYLGTIRNYEFRLNTAVHRAERPTFPERAHKLGYKAKQGICIFRVRIRRGGRKRLVNNGNTRGKPVNAGIYQLKPANSLQSMAELKAGKRAGNLRVLNSYWVGQDGVYKYFEVIMVDPNHNAIRNDPKLNWICKSTMKHRECRGLTSASRKSRGLGKGIRYNHTIGGSRKAAWRRRNTVSLRRYR |
| LP0 | uL22 | MKKKYAPEIHDPGNTVKCRIDNARVSFKNTRETSRVLVNRKLGDALNYLYDVIKKKQCVPMKRYARGVGRTAQAKAFKTQRGRWPVKSAKFFIELINNLKVNAVNKNLNPDEMVIKNIIVNKAPIIPGRIHRAYGRINPYNSHPCHIQMIAVKKMVAVPKAIDGEYAAEVV |
| LQ0 | eL18 | MVATAFASKKYVSRGRKKLVSRNIYLEGLAALYERIAKSSANEIVHKIAAQLKMSRNNRPPMKVSKLANISEKYPGKVYVVVAKVLDDNCFLEVPKMQIVALQFSKSAKEKIEKAGGSTHTLDQLFEVAPGLENVELFRGDLTARKAYKYFGAPGDRHSRTYPKTTSKGKNREKRLLK |
| LS0 | eL20 | MWRCGGIKEYRIYGSKMPTEQEVAPQIFFHNVFAKNEIVARSRFNNLMKTKYKIKSGKIVILKIEELVEDLKDMKIKNYGIQLVYRSKKGIHNMYKEFRSISRCKAVEMLFNDMAGRHKAKRDDIKIVSLKELRSEDLRRDRVIQFTKDDVMYPVFKKVLNSKYDFVLKGTNIFN |
| LR0 | eL19 | MTKLPTIRRIAASIFKCGKNKIWFDPGETPRLASSSTRMQVRRLIKDGVILRKPNTVHSRWRANKRAEARKKGRHMGIGKRKGTKNARMPEKRVWIKKIRGQRASLKEMKKSGHITPEEFRQYYMQAKGNMFKSLKVMEDHIEKKREEKMRIKDLAAQAAALRMKK |
| LT0 | eL21 | MRSNGYRRGTRHRFSQAFRKHGMPKPSILTRVFKKGQYVDVVVNPAIHKGMPHKFFHGRTGKIFNIDKRSIGILMNKRCGPRYVEKMVIARVEHVRPSRCNEEYIKRRTENDRLRREAAARGEKLGSLKRKPGGPRGAVLISTENNTPIEIRNEPYFEVY |
| LU0 | eL22 | MNSVQAENDTRCYELNCSVLVKDSLLSTEDLNSYLEARMKVNGRTGNLAGNIDLVCTEESILVKPRVRLSKKYLKYLVKKFLYKKELIDWVRILSTGKSSYKLAYYRVVSNKYE |
| LV0 | uL14 | MPSKHVLEDRRPKIRSTCGVQVGTRIACADNTGAKIMQIIGVKGVRGRLNRLPAASVGDVVLCSVKKGRPDMRKKIVLCVVIRQKKAWRRRDGSHICFEDNAAVVITNKGDPKGTQIAGPVPREVADIWPKISSNAPAII |
| LX0 | uL23 | MEIKRNNPKYTRKAVTQPATGHPADIIRFGACNEKAVRLIENNNTLVFICDKYATKPQIGNAVTRFYKVPVEKVNTARSIKGYKKAYVKLKNEGDALKIANEAGII |
| LW0 | eL24 | MVKEGVCVYSGYEVPKGSGLIRVTNDTRSFLFLNKKVQSLSNRKINPRDVAWTAASRAFHKKGMKKVVRKEAEIQVVKEVRGFPSVPKSIIVQPKRDDDQKKKAERGAETFRAQKKVTKAEGRKMKADGRWQR |
| LY0 | uL24 | MKFNKEKTASRRKNRKAHFTANSTERRIRMSSPLSKELREKYGFRSFPIRRNDKVVVMKGRFKGNTGTVVEVRRKMYKVYVDSCEASKMNGRKVRVGIDASNLKIIELYQGDGRDKILERKMNNRRMQMERAAKIKMEQK |
| LZ0 | eL27 | MLLKPNTVVVALKGRFAGKKGVVVSASEDGRKILVAGIEKMPQPVTDDMSECKKKRLSRMSAFIKNYNSRHLLATRYYGDVGLGAIDFSRIFENFESKKMAIDAVKKAFINANKQKKAAWLFEKLVF |
| LAA | uL15 | MYQNFYPMASKDKKTRKLRGHVSHGHGRVGKHRKHPGGRGKCGGMAHRKTLFMKYHPDHFGKRGMNCTHLKKNARYAPPINVSKLWSLIPKSQLETIMNDNTIAPIINCRSFGYHIVRGGGQLSLKRPIVVMARYFTPKAVSMIESLGGRCIISP |
| LBB | eL29 | MAKRKNHTNHNQNRKNHRNGIKKVKKSAPSFRGLNHKYLRNMLYSRKYNNIGRAAYEAEHGPQQ |
| LCC | eL30 | MSRKRNVTDGLAFKLPLAMRTGVYKVGYKSAIKLLQAGRTKYIVAAANFPSVKRKLLEYYAALSNNVPVVIFKGSNNELAKVCDHHYRIGVISILDDGESGLISAGTQ |
| LDD | eL31 | MASLSNKAVEMTVNLGKLARKASWRYKAPKCIYYLKKFIRSQFKSENDILIAPEVNKYIWRHGIKNIPKRMRIKIERGPSNKNPELNVFRVCLVNVNTFKGLQSQSYTE |
| LEE | eL32 | MEPYLRVPRQRKTKHKFNRFHSDRFKRVKKSWRRPKGIDNRVRRKLSGAIKMPNKGYGTDKLAKHMVHGGFRLVQIRNVGDLLPLISQNRFYCAEVAHSVGAKKRIEIFYKAMEYGIHLTNGKARLVEENKE |
| LGG | eL34 | MVQHLTHRGRRTYNNASNQKKIRRTPGNRRVYIPKKKPGRVPKCVKCRSKLRGIDICRPAAFARLRKSQRTVARTYGGNLCGSCLENRILDAFLSAEEQLLLGKKQTNAADNN |
| LHH | uL29 | MKIRTLELREKSVEELETLAFSLKEELLRLRQQKNTQTLKPHEIRVMRKNIARVLTVRTEKLYAEMYEKHKNDKRMPKDLRPRLTKAKRMALTKRQLNMSVHSVCKRRRAYPRMYFSYSEPVDPKN |
| LFF | eL33 | MKDVFDPCLTAIPATFISHMRGRRIVHPSHSLLRIEGVKTREDAKFFLGNGVVSLYELSGKKMENRGRIRRLHGNSGVVAAKFERNLPPNKIGTEVFVKLYKVEDDDY |
| LII | eL36 | MRKKLRRMRVKYPVTPLVGIEPIPRPGKKHDWNNHGSPMAKAIASEICGLAPYEKKALELIKSDQERKCRRFLKKRLGNLTRTKRKQEALTAIAREQQ |
| LOO | eL42 | NGKWQNNLKVNIPDKRNTHCKKCNKHTEHKVSQYKKSKEGRNGQGTRRYRRKQRGYHGQTKPILRRKAKVTKKLVIKLRCSVCDAKHQQTMKRTKHVEFGGKKKTKGQALTY |
| LJJ | eL37 | SKGTSSFGKKNKRNHILCVRCGSMSYHKQKLRCSSCGYPEKKLRNRGSPKARRRRGEGTGRMRHLKKVRRAARNGFKGNAILRALRNANTTSDSK |
| LPP | eL43 | AKSRGTKKVGIVGKYGVRYGSSLRKRIKAIEISQHAKYECKACGKTSVKREVVGIWKCKACAFTFAGGAFAPTTSAGAHSNSITRQ |
| LLL | eL39 | GSRKTEIIKTRLSRALVRNRPIPVWKRRMRGNTQQYNMKRRHWRRNKLKIY |
| LMM | eL40 | MQVFIKSPGLLEVVSIDRDTTISDLQAMSRYLCNMAFVVNGVLLDKDISLASQGIQDLSTVSAIPLLVGGAGDKQMNENDKALALKRKEAMICRSCYARLAPRAINCRKKKKCGGSNNLRPKKKLKETKKGK- |
